# Cortical thinning in temporal pole, a core region in Alzheimer’s disease, in non-demented, middle-aged *APOE*-ε4 and *PICALM*-AA/AG carriers

**DOI:** 10.1101/2025.02.21.639542

**Authors:** Patrycja Dzianok, Jakub Wojciechowski, Tomasz Wolak, Ewa Kublik

## Abstract

The symptoms of Alzheimer’s disease (AD) are caused by neurodegeneration and atrophy in particular brain regions, especially in the temporal cortex. However, the influence of genetic risk on cortical thickness in non-demented individuals prior to disease onset remains unclear. This study aimed to explore the relationship between two AD risk genes (*APOE/PICALM*) and cortical thickness in selected regions of interest (ROIs) in non-demented, middle-aged individuals. Sixty-nine (N = 69) participants (34 females, 35 males; age: 55.45±3.19) underwent magnetic resonance imaging (MRI). They were divided into three groups based on their AD risk. Cortical thickness was analyzed using CAT12 software (surface-based morphometry with the Destrieux atlas) based on T1-weighted MR images in five ROIs referred as “the cortical signature of AD” in previous studies. *APOE*-ε4 with *PICALM*-AA/AG carriers (A+P-) are characterized by a thinner cortex in the right temporal pole compared to non-carriers, controlling for sex. No other differences in cortical thickness were found in the selected ROIs. The direction of the findings aligns with existing literature reporting cortical thinning in amyloid-positive individuals, as well as in patients with mild cognitive impairment and Alzheimer’s disease when compared to control groups.

## Introduction

Genetic factors play a pivotal role in the pathogenesis of late-onset Alzheimer’s disease (LOAD), with several genes identified as significant contributors to the risk of developing the disease. Among these, the apolipoprotein E (*APOE*) has emerged as a key genetic factor (Michaelson 2014). The *APOE*-ε4 allele, in particular, has been consistently associated with increased Alzheimer’s disease (AD) risk and earlier onset of the disease (Corder et al. 1993; Saunders et al. 1993). *PICALM*, being one of the most influential AD risk-genes (Jun et al. 2010; Lambert et al. 2011; Ando et al. 2022), has also been implicated in the modulation of AD pathology through its involvement in clathrin-mediated endocytosis, which is critical for amyloid precursor protein processing and amyloid-β (Aβ) clearance (Xu et al. 2015). Several *PICALM* single nucleotide polymorphisms (SNPs) are associated with AD, but the PICALM rs3851179 variant is the most studied, with the G/G alleles considered a risk factor. Both *APOE* and *PICALM* are implicated in Aβ pathology (Xu et al. 2015; Robinson et al. 2018), and a genetic interaction between these two genes that increases AD risk has been identified. This interaction has been observed in meta-analyses involving diverse populations, including White, African American, Israeli-Arab, Caribbean Hispanic (Jun et al. 2010), and Chinese cohorts (Chen et al. 2012). According to those studies, *PICALM* is considered a risk factor only in *APOE*-ε4 carriers.

Cortical thickness, which refers to the measurement of the outer layer of the brain known as the cortex, is widely recognized as a sensitive indicator of neurodegeneration (Singh et al. 2006; Bakkour et al. 2009; Dickerson et al. 2009; Schwarz et al. 2016), as gray matter atrophy is a hallmark feature of Alzheimer’s disease. This parameter is also of considerable interest in studies of other neurological disorders and in the context of normal brain aging (Shaw et al. 2016). While gray matter loss can be estimated using volumetric analysis, cortical volume is less preferred in Alzheimer’s disease biomarker research due to potential confounding factors, such as variations in head size (Schwarz et al. 2016). Nonetheless, both cortical thickness measurement and volumetric analysis demonstrate comparable effectiveness in differentiating Alzheimer’s disease patients from healthy controls (Schwarz et al. 2016).

Cortical thinning in dementia is often region-specific (Gutiérrez-Galve et al. 2009) and typically progresses as the disease advances (Singh et al. 2006; Schwarz et al. 2016). In 2019, nine regions were identified as the “cortical signature of Alzheimer’s disease” with five regions—the parahippocampal gyrus/medial temporal cortex, supramarginal gyrus, temporal pole, superior parietal lobule, and inferior temporal gyrus—highlighted as critical for tracking the progression from subjective cognitive decline in healthy individuals to Alzheimer’s disease (Bakkour et al. 2009; Dickerson et al. 2009; Verfaillie et al. 2016). Notably, cortical thinning frequently begins before the onset of clinical symptoms, suggesting its potential as an early biomarker (Bakkour et al. 2009; Dickerson et al. 2009).

Studies have also indicated that genetic factors, including *APOE* (Burggren et al. 2008; Espeseth et al. 2008; Donix et al. 2010; Donix et al. 2013) and *PICALM* (Wu et al. 2022), may independently influence cortical thickness across the lifespan in healthy individuals. The interaction between *APOE* and *PICALM* has been shown to influence brain atrophy in Alzheimer’s disease patients (Morgen et al. 2014), as well as impact memory and cognitive performance (Morgen et al. 2014; Chang et al. 2019). Additionally, it has been associated with a reduced cerebral metabolic rate for glucose in the default mode network in these patients (Chang et al. 2019). However, the impact of *APOE-PICALM* interaction on cortical thickness in non-demented adults remains unexplored.

Despite extensive research into the genetic underpinnings of Alzheimer’s disease, the mechanisms by which polymorphisms in genes such as *APOE* and *PICALM* jointly influence brain structure remain poorly understood. In this study, we aimed to examine the impact of *APOE* and *PICALM* polymorphisms on cortical thickness in middle-aged individuals at genetic risk for Alzheimer’s disease who have not yet exhibited symptoms of cognitive decline. These findings could offer valuable insights into the early neurobiological changes associated with Alzheimer’s disease risk.

## Material and methods

### Study design, participants, and study groups

We established the PEARL-Neuro Database (Dzianok and Kublik 2024), detailing the study methodology, including comprehensive information on study inclusion/exclusion criteria. Briefly, the screening phase involved N = 200 middle-aged participants (aged 50-63). Genetic tests (Sanger sequencing) were conducted to obtain information on two AD risk genes: apolipoprotein E (*APOE*) and phosphatidylinositol-binding clathrin-assembly protein (*PICALM*). *APOE*-ε3/ε4 and *APOE*-ε4/ε4 were considered risk variants, while *APOE*-ε3/ε3 was considered a neutral variant; no participants with the ε2 allele were included in the analysis. For *PICALM, PICALM*-G/G was considered a risk variant, and *PICALM*-A/A and *PICALM*-A/G were considered neutral variants. In the second phase, MRI scans were performed on N = 69 participants (The neuroimaging phase of the PEARL-Neuro Database originally consisted of 79 participants. However, not all participants ultimately met the inclusion criteria for MRI, and some withdrew from this part of the study, which is described in full in Data Note about this dataset (Dzianok and Kublik 2024)). Finally, participants were categorized into three groups based on varying levels of LOAD risk: N (neutral variants of *APOE/PICALM*) group: N = 27; A+P-(risk variants of *APOE* and neutral variants of *PICALM)*: N = 24; A+P+ (risk variants of *APOE/PICALM*): N = 18.

The study was approved by the local bioethics committee (Bioethics Committee of the Nicolaus Copernicus University in Toruń at Collegium Medicum in Bydgoszcz, Poland; ID of the approval: KB 684/2019). All participants provided written informed consent for study participation and received financial compensation.

### Statistics

Firstly, the linearity between age and all variables was checked. Since no linearity was found, group differences were assessed using a two-way ANOVA with sex as an additional factor. If the assumptions for two-way ANOVA (normality via Shapiro-Wilk test and QQ plots, factor normality, and homogeneity of variances via Levene’s test) were not met, the assumptions for one-way ANOVA were checked instead. If the assumptions for one-way ANOVA were also not met, the non-parametric Kruskal-Wallis test was used. The specific test used for each analysis is indicated in the results tables.

### Data visualization

Boxplots were generated using custom Python scripts, with use of the seaborn (Waskom 2021), matplotlib (Hunter 2007), and statannot (Weber 2022) libraries. The brain figure illustrating the ROIs was created using BrainPainter with Destrieux atlas data (Marinescu et al. 2019).

### MRI

T1-weighted images were acquired at the end of the neuroimaging session conducted at the Bioimaging Research Center, Institute of Physiology and Pathology of Hearing, Poland. Imaging was performed using a Siemens Prisma FIT 3T scanner (Siemens Medical Systems, Erlangen, Germany) equipped with a 64-channel phased-array RF head coil. The acquisition parameters of T1-weighted 3D MP-Rage images were as follows: TR = 2400 ms; TI = 1000 ms; TE = 2.74 ms; flip angle = 8 degrees; FOV = 256×256 mm; voxel size = 0.8×0.8×0.8 mm; TA = 6:52 min.

### Cortical thickness analysis

We used Computational Anatomy Toolbox (CAT12) (Gaser et al. 2022) to analyze cortical thickness (using Destrieux atlas (Destrieux et al. 2010)) with a surface-based morphometry approach, specifically the projection-based thickness method proposed by Dahnke and colleagues (Dahnke et al. 2013).

We focused on five specifically chosen regions of interest (ROIs) identified as indicative of progression from subjective cognitive decline in healthy individuals to Alzheimer’s disease (Verfaillie et al. 2016), which were based on nine AD-related areas named “the cortical signature of AD” (Bakkour et al. 2009; Dickerson et al. 2009). The chosen ROIs thickness was compared between the study groups and included the parahippocampal gyrus (a component of the medial temporal cortex), supramarginal gyrus, temporal pole, superior parietal lobule, and inferior temporal gyrus.

## Results

### Cohort characteristics

The MRI study cohort consisted of right-handed, non-demented, middle-aged (age: 55.45±3.19) adults, with an equal distribution of females and males (34 females, 35 males). In a related article (Dzianok et al. 2024) from our PEARL-Neuro Database (Dzianok and Kublik 2024), we have previously reported no differences among the study groups regarding age, education, gender, handedness, potential alcohol issues, smoking status, and overall health status. Additionally, the information about psychological test assessments and other health-related AD risk factors is also summarized in that paper (Dzianok et al. 2024).

### Cortical thickness

A+P-group was characterized by a thinner cortex in the right temporal pole than the N group (F(2,63) = 3.59, *p* < .05, post-hoc: *p* < .05; Figure 1, Figure 2, Table 1). The groups did not show any differences in cortical thickness in the other analyzed ROIs.

**Table 1.** Descriptive statistics and group differences (*Group* factor) in cortical thickness.

| Area | Group |  |  | p-value |
| --- | --- | --- | --- | --- |
|  | A+P+ | A+P- | N |  |
| Parahippocampal gyrus left | 2.81 ± 0.25 | 2.78 ± 0.18 | 2.78 ± 0.22 | .81 |
| Parahippocampal gyrus right | 2.88 ± 0.20 | 2.90 ± 0.16 | 2.88 ± 0.19 | .93 |
| Supramarginal gyrus left | 2.56 ± 0.13 | 2.53 ± 0.11 | 2.53 ± 0.12 | .73 |
| Supramarginal gyrus right | 2.57 ± 0.14 | 2.54 ± 0.10 | 2.57 ± 0.12 | .69 |
| Temporal pole left | 3.14 ± 0.24 | 3.07 ± 0.18 | 3.13 ± 0.20 | .42 |
| **Temporal pole right** | **3.19 ± 0.24** | **3.07 ± 0.12** | **3.19 ± 0.19** | **< .05** |
| Superior parietal lobule left | 2.56 ± 0.13 | 2.53 ± 0.11 | 2.53 ± 0.12 | .58 & |
| Superior parietal lobule right | 2.57 ± 0.14 | 2.54 ± 0.10 | 2.57 ± 0.12 | .64 & |
| Inferior temporal gyrus left | 2.67 ± 0.15 | 2.67 ± 0.14 | 2.69 ± 0.15 | .91 |
| Inferior temporal gyrus right | 2.71 ± 0.20 | 2.70 ± 0.17 | 2.70 ± 0.17 | .97 |

<sup>a</sup>& Kruskal-Wallis test. The bolded font indicates a significant result. Descriptive statistics: mean ± standard deviation.

**Figure 1.**
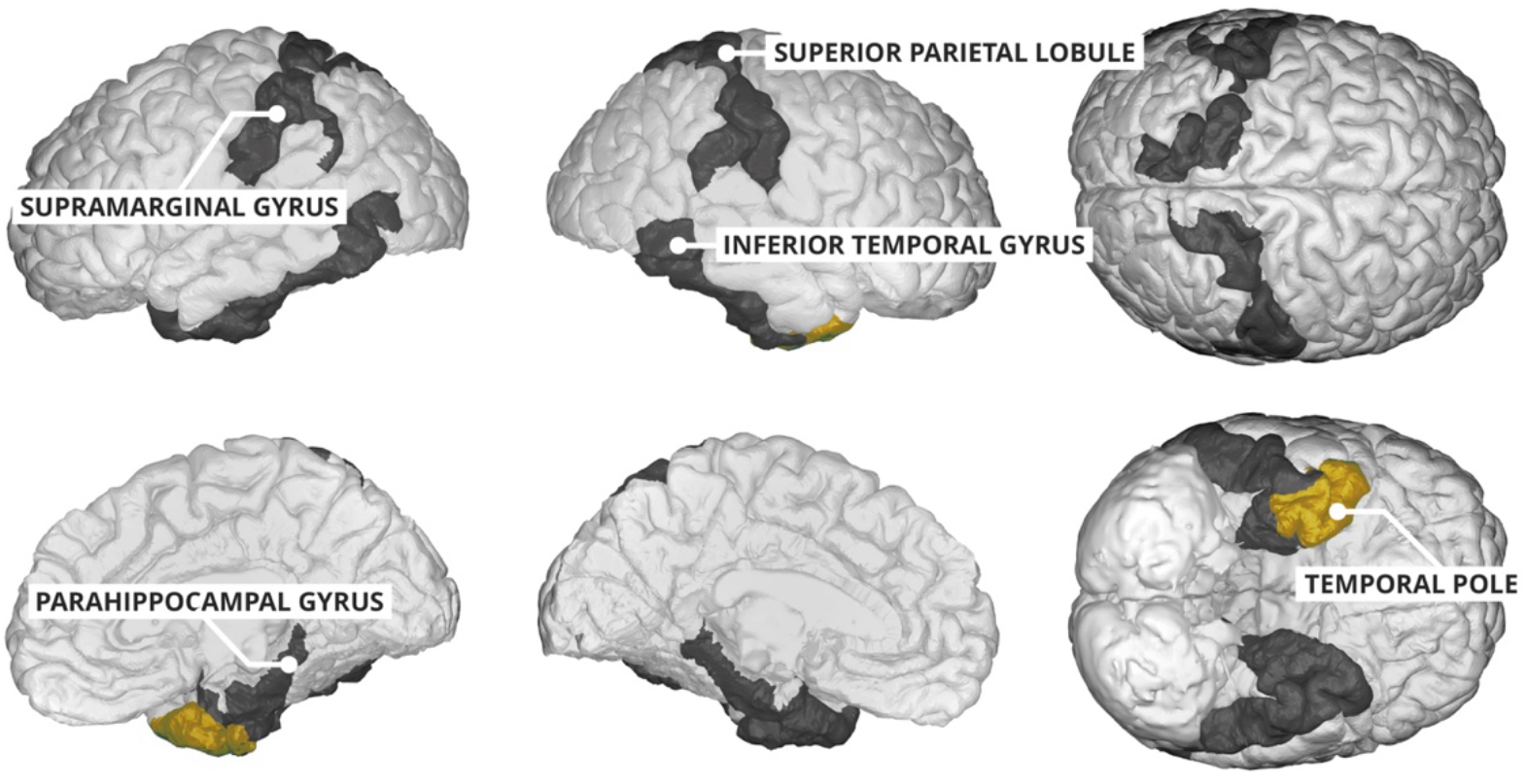
The analyzed ROIs are marked in black. The ROI (the right temporal pole) that was significantly thinner in the A+P-group compared to the N group is highlighted in yellow.

**Figure 2.**
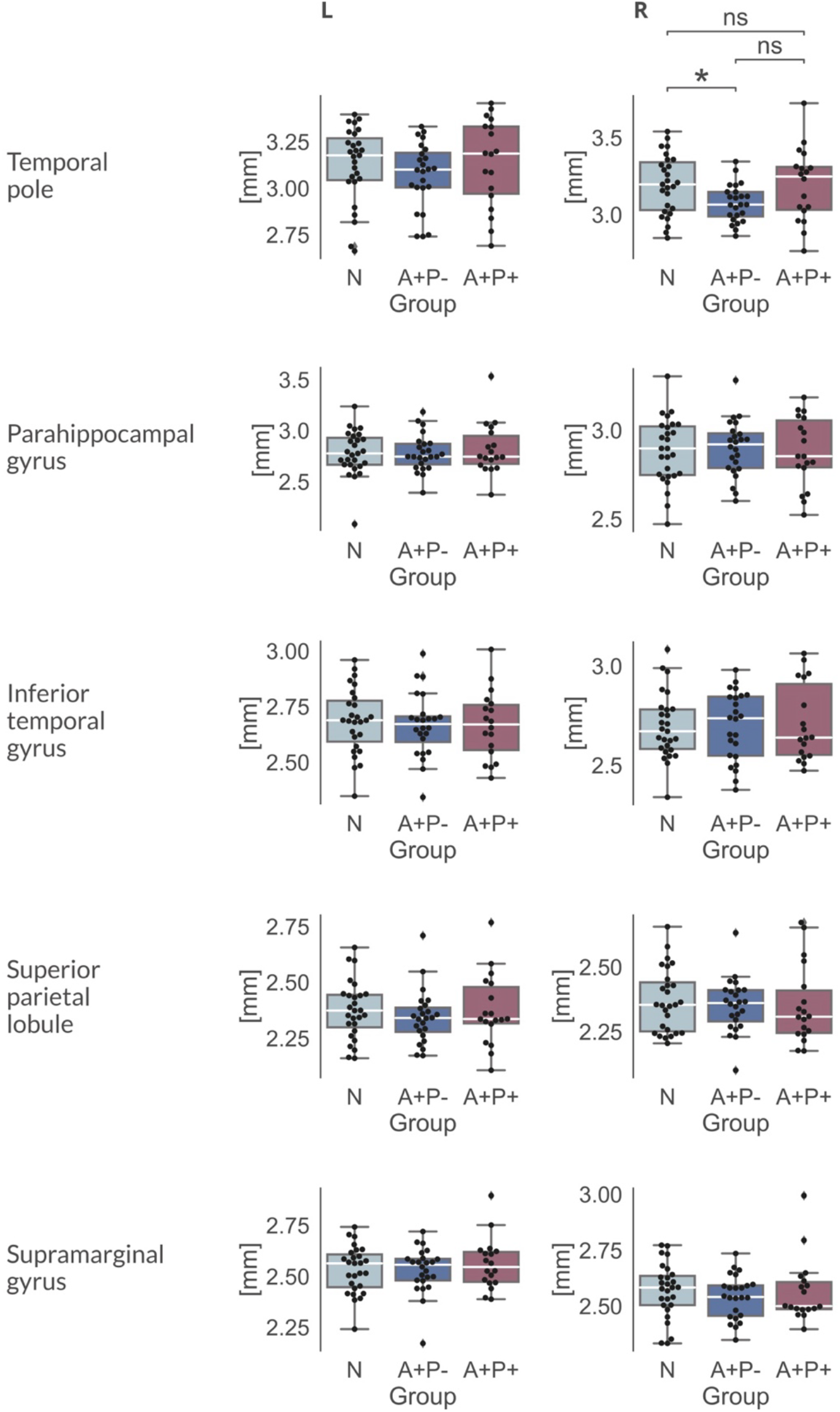
Group differences in cortical thickness (in mm) across all analyzed ROIs are presented. Significance between the groups is only indicated for the ROI marked as significant in Table 1, for clarity. * – *p* < .05, ns – not significant. No differences were observed between the groups in the other ROIs.

Additionally, females tended to have thicker cortex at left temporal pole than males (F(1,63) = 8.98, *p* < .01; Table 2), and similar trend was observable for right temporal pole (F(1,63) = 3.39, *p* = .07) and left supramarginal gyrus (F(1,63) = 3.47, *p* = .07; Table 2).

**Table 2.** Descriptive statistics for the *Sex* factor, along with the significance for differences associated with the *Sex* factor and the interaction effect (*Sex*Group*). The results are shown only for the ROIs that met all the assumptions for two-way ANOVA, as outlined in the Methods section.

| Parameter | Sex |  | p-value Sex | p-value Sex*Group |
| --- | --- | --- | --- | --- |
|  | Females | Males |  |  |
| Parahippocampal gyrus left | 2.80 ± 0.21 | 2.79 ± 0.23 | .85 | .22 |
| Parahippocampal gyrus right | 2.89 ± 0.17 | 2.88 ± 0.18 | .83 | .46 |
| Supramarginal gyrus left | 2.57 ± 0.11 | 2.51 ± 0.12 | .07 | .59 |
| Supramarginal gyrus right | 2.56 ± 0.12 | 2.56 ± 0.12 | .94 | .30 |
| **Temporal pole left** | **3.18 ± 0.17** | **3.04 ± 0.22** | **< .01** | .28 |
| Temporal pole right | 3.19 ± 0.20 | 3.11 ± 0.18 | .07 | .11 |
| Inferior temporal gyrus left | 2.69 ± 0.14 | 2.66 ± 0.15 | .48 | .41 |
| Inferior temporal gyrus right | 2.72 ± 0.17 | 2.68 ± 0.18 | .43 | .89 |

<sup>a</sup>The bolded font indicates a significant result. Descriptive statistics: mean ± standard deviation.

## Discussion

We demonstrated that *APOE* risk carriers with *PICALM* neutral alleles (A+P-group) exhibit a thinner cortex in the right temporal pole compared to non-risk carriers (N group). On average, this difference was approximately 0.12 mm, which is about half the typical difference (∼0.25 mm) observed between individuals with Alzheimer’s disease and healthy individuals (Dickerson et al. 2009). There were no significant differences in cortical thickness between the groups in the parahippocampal gyrus, inferior temporal gyrus, superior parietal lobule, or supramarginal gyrus—regions that are also implicated as key cortical areas in AD and are involved in the progression from subjective cognitive decline to AD (Bakkour et al. 2009; Dickerson et al. 2009; Verfaillie et al. 2016). Consistent subtle thinning was previously observed in some of the above-mentioned ROIs among amyloid-positive asymptomatic individuals compared to amyloid-negative controls, with this effect being particularly evident in the temporal pole (Dickerson et al. 2009). Another study, using data from the Alzheimer’s Disease Neuroimaging Initiative (ADNI), showed increased cortical thickness loss predominantly in the left temporal pole (anterior part) among patients with mild cognitive impairment who converted to Alzheimer’s disease (Iannopollo et al. 2021). Other early research from the 1980s and 1990s also suggests that the temporal pole is one of the regions affected early in the course of AD (Van Hoesen et al. 1986; Arnold et al. 1991; Arnold et al. 1994). Alzheimer’s disease patients exhibit significant atrophy in this area, with primary neuronal loss observed in layers III and V, and neurofibrillary tangles identified in layers II, III, V, and VI (Iannopollo et al. 2021). It was previously demonstrated that cortical thinning, particularly in the temporal poles and left medial temporal lobe, occurs in healthy adults who later developed cognitive impairment (diagnosed as MCI or AD) compared to those who remain cognitively healthy (Pacheco et al. 2015). All this evidence, along with our new findings on non-demented individuals, suggests that the thickness of the temporal pole may be the first subtle, yet consistent marker associated with various well-documented AD risk factors (genetic risk in this case), and the development of Alzheimer’s disease (as documented in other studies).

Two general hypotheses have been proposed to explain how genes, particularly *APOE*, may influence healthy adults, and these mechanisms are not mutually exclusive (O’Donoghue et al. 2018). The first is the prodromal hypothesis, which states that the observed genetic impact may reflect the earliest stages of undiagnosed dementia. The second is the phenotype hypothesis, which suggests that genes, such as *APOE*, contribute directly to variations in brain function and cognition, potentially predisposing individuals to develop dementia. Reduced cortical thickness in the medial temporal lobe regions and hippocampus has been previously reported in non-demented *APOE*-ε4 carriers (Burggren et al. 2008; Donix et al. 2010; Donix et al. 2013). Hemispheric asymmetry associated with *APOE*-ε4 has also been noted in healthy controls as well as in individuals with MCI or AD, supporting our finding that the effect is predominantly observed on one side—specifically the right side. Some other studies have identified greater cortical thinning associated with *APOE* in the left hemisphere (Donix et al. 2013), while others have reported greater thinning in the right hemisphere (Goñi et al. 2013). Additionally, case studies have documented highly asymmetric amyloid-beta (Aβ) deposition and neurodegeneration predominantly affecting the right hemisphere (Lee et al. 2021). Conversely, other research indicates that local gray matter loss rates in Alzheimer’s disease are more pronounced in the left hemisphere (Thompson et al. 2003). While most studies report *APOE*-ε4-related cortical thinning, one study found that middle-aged *APOE*-ε4 carriers (aged 48 and above) exhibited thicker cortex in several regions earlier in life, followed by a steeper age-related cortical decline in areas such as the temporal poles (Espeseth et al. 2008). These findings align with the antagonistic pleiotropy hypothesis (Tuminello and Han 2011; Rusted and Carare 2015; O’Donoghue et al. 2018), which posits that a single genetic variant may exert beneficial effects on certain traits during early life while contributing to negative outcomes in aging.

Limited data exist on the impact of *PICALM* on brain anatomy in healthy participants. A recent study (Wu et al. 2022) reported that PICALM rs3851179 similarly modulates cortical thickness and cerebrospinal fluid (CSF) biomarkers in both Alzheimer’s disease patients and healthy elderly controls. Interestingly, the study found that levels of Aβ42 (and the Aβ42/40 ratio) were higher, phosphorylated and total tau (P/T-tau) levels were lower, and cortical thickness in the postcentral area was reduced in individuals with the *PICALM*-AA/AG genotype (considered neutral) compared to those carrying the risk-associated G allele. Our findings appear to align with this pattern: *APOE*-ε4 carriers with the *PICALM*-A genotype (A+P-group) exhibited thinner cortex in the temporal pole compared to control participants. Conversely, in the same study (Wu et al. 2022), AD patients with the *PICALM*-A genotype, despite demonstrating greater Aβ accumulation, performed better on the Mini-Mental State Examination (MMSE), suggesting a complex interplay between *APOE, PICALM*, and AD development.

Previous studies have demonstrated that the interaction between *APOE* and *PICALM* has a negative impact on brain atrophy in Alzheimer’s disease patients, particularly contributing to reduced volume in the prefrontal regions (Morgen et al. 2014). However, the combined effects of these genes on brain atrophy in non-demented individuals remain poorly understood. In this study, we observed that cortical thickness is reduced in single-risk *APOE* carriers (with neutral *PICALM* alleles: A/A and A/G) but not in the ‘double-risk’ group carrying both *APOE* and *PICALM* risk variants. This finding contrasts with our initial hypothesis, which anticipated that the observed effects would increase proportionally with the cumulative genetic risk. Instead, our results align with the previously mentioned study (Wu et al. 2022), suggesting that the neutral A allele of the *PICALM* gene is associated with reduced cortical thickness—in this study particularly in the left postcentral cortex.

In this study, we did not examine any amyloid-beta (Aβ) markers, such as those found in blood or cerebrospinal fluid; therefore, we cannot directly determine whether this kind of AD-related pathology had begun to develop in our participants. However, it is well-established that individuals carrying the *APOE*-ε4 allele are more prone to Aβ accumulation (Morishima-Kawashima et al. 2000; Lim et al. 2017; Baek et al. 2020), a key process leading to cortical neurodegeneration.

No interaction between the genetic *Group* and *Sex* factors was identified in our analysis. However, the inclusion of the *Sex* factor revealed that females exhibited thicker cortical regions in the left temporal pole, with a similar trend observed in the right temporal pole and the left supramarginal gyrus. Previous studies have reported gender differences in cortical thickness, such as thicker cortices in women, including temporal regions (Sowell et al. 2007; Ritchie et al. 2018). As our primary focus was to use the *Sex* factor as a covariate to control for its influence on our main hypothesis, we will not discuss these findings in detail.

Future longitudinal studies, examining the same individuals over time, are planned to further investigate questions arising from this phase of the research.

### Limitations

An important methodological consideration in this study involves the representation of selected brain regions. Structural neuroimaging software provides access to various brain atlases for region selection. To test the hypothesis concerning cortical thinning, we employed a surface-based morphometry approach. To enhance the statistical power of our study, we selected regions of interest using the Destrieux Atlas, one of the most widely used atlases. Its standardized anatomical nomenclature and broad adoption in the literature facilitate comparisons with existing studies (Destrieux et al. 2010). However, it is important to acknowledge that the regions defined by the Destrieux Atlas may not correspond precisely to those used in studies employing voxel-based morphometry (VBM) or vertex-wise cortical mapping methodologies. For instance, while Dickerson’s study highlights the medial temporal cortex (MTC) as a significant area (Dickerson et al. 2009), the Destrieux Atlas delineates individual sulci and gyri rather than broader cortical regions. In our study, the medial temporal cortex is represented by the parahippocampal gyrus, a critical component of the MTC, located around the hippocampus and essential for memory encoding and retrieval. This methodological distinction highlights the importance of cautious interpretation and careful cross-study comparisons, particularly when different atlases or mapping approaches are employed.

## Acknowledgements

We would like to thank the participants for their involvement in the study.

## Data availability statement

A neuroimaging dataset and information about participants are available in open-access dataset PEARL-Neuro (Dzianok and Kublik 2024). Structural MRI data are protected and available upon request. To access these data, please contact us. Access is granted for research purposes under a Data Use Agreement (DUA).

## Authors’ contributions

PD: Conceptualization; Data curation; Formal analysis; Funding acquisition; Investigation; Methodology; Project administration; Software; Visualization; Writing – original draft. JW: Formal analysis; Methodology; Software; Writing – review & editing. TW: Methodology; Resources; Supervision; Writing – review & editing. EK: Conceptualization, Supervision; Writing – review & editing.

